# Can net diversification rates account for spatial patterns of species richness?

**DOI:** 10.1101/240978

**Authors:** Camilo Sanín, Iván Jiménez, Jon Fjeldså, Carsten Rahbek, Carlos Daniel Cadena

## Abstract

The diversification rate hypothesis (DRH) proposes that spatial patterns of species richness result from spatial variation in net diversification rates. We developed an approach using a time-calibrated phylogeny and distributional data to estimate the maximum explanatory power of the DRH, over a given time period, to current species richness in an area. We used this approach to study species richness patterns of a large family of suboscine birds across South America. The maximum explanatory power of the DRH increased with the duration of the time period considered and grain size; it ranged from 13 – 37 fold local increases in species richness for *T* = 33 Ma to less than 2-fold increases for *T ≤* 10 Ma. For large grain sizes (≤ 8° × 8°) diversification rate over the last 10 Ma could account for all the spatial variance in species richness, but for smaller grain sizes commonly used in biogeographical studies (1° × 1°), it could only explain < 16% of this variance. Thus, diversification since the Late Miocene, often thought to be a major determinant of Neotropical diversity, had a limited imprint on spatial richness patterns at small grain sizes. Further application of our approach will help determine the role of the DRH in explaining current spatial patterns of species richness.

**Note to readers:** This manuscript has been seen by a few researchers, some of whom suggested that before publishing our work in a peer-reviewed journal we should conduct simulations to demonstrate that our methods properly estimate the contribution of variance in diversification rates to spatial variation in species richness. Although we believe that our approach derives logically from theory and statistics and is therefore valid, we understand that it is rather unique and see why some readers would think that an independent validation is necessary. Unable to complete such validation in the near future, however, we decided to make this manuscript available as a preprint to share our ideas and hopefully stimulate discussion on what we believe is a most interesting topic. We also hope to receive feedback that may enable us to improve our work for publication in a journal at a later date.

## INTRODUCTION

Broad-scale spatial patterns of variation in biological diversity are well documented for several taxa (Hawkins et al. 2003; Rahbek and Graves 2001; Whittaker et al. 2001; Willig et al. 2003), but it is unclear how these patterns arise from the three fundamental processes determining species richness: speciation, immigration, and extinction (Jablonski et al. 2006; Ricklefs 2006a; Rosenzweig 1995; Wiens 2011; Wiens and Donoghue 2004). The Diversification Rate Hypothesis (DRH; Fischer 1960; Mittelbach et al. 2007; Ricklefs 2004) proposes that geographical patterns of species richness result from spatial variation in net diversification rate, the balance between speciation and extinction rates. According to this hypothesis, current species richness is high in areas experiencing high speciation rates and/or low extinction rates. Additional hypotheses proposed to explain geographical patterns of species richness incorporate diversification rate (Kisel et al. 2011; Svenning et al. 2008), including the “cradle” and “museum” hypotheses (Chown and Gaston 2000) as well as hypotheses positing that ecological limits to speciation (Rabosky 2013; Rabosky and Hurlbert 2015), energy and water availability (Currie et al. 2004; Hurlbert and Stegen 2014), topographic heterogeneity (Fjeldså 1994; Rahbek and Graves 2001), biotic interactions (Schemske 2009), or climatic oscillations (Dynesius and Jansson 2000; Fjeldsâ et al. 2012) affect net diversification rate leaving an imprint on contemporary spatial patterns of richness.

Although previous studies (e.g. Cardillo et al. 2005; Jansson et al. 2013; Jetz et al. 2012; Pyron 2014; Pyron and Wiens 2013; Ricklefs 2006c; Rolland et al. 2014; Soria-Carrasco and Castresana 2012; Svenning et al. 2008) tested key predictions of the DRH and related hypotheses (e.g., that areas with high species richness harbor clades that have experienced relatively high rates of net diversification), it appears that two crucial quantitative questions have not been addressed: (1) what is the contribution of net diversification over a given time period to current species richness in an area? and (2) how much of the spatial variance in current species richness can be explained by the spatial pattern of net diversification rate? Answering these two questions amounts to evaluating the explanatory power of the DRH. Doing so is useful because spatial patterns of species richness are thought to be best explained by a combination of several non-exclusive hypotheses that emphasize different processes (Brown 2014), and further understanding requires establishing the extent to which each hypothesis can account for different facets of these patterns. Indeed, the explanatory power of any given hypothesis may depend on the temporal and spatial scales considered (Fine and Ree 2006; Rahbek and Graves 2001), as we describe below in relation to the DRH.

Because the contribution of spatial variation in net diversification rates to patterns of species richness may vary temporally, the answer to the two quantitative questions above may depend on the time window considered. For instance, the diversification rate of a clade could decrease over time if clade diversity were regulated by ecological factors (Rabosky 2009; Rabosky et al. 2012); if these limiting factors were structured geographically, then current spatial patterns of species richness could be determined more strongly by early diversification than by recent diversification. Likewise, current richness patterns of palms (Svenning et al. 2008) and birds (Fjeldså and Rahbek 2006; Thomas et al. 2008) in South America could partly reflect spatial variation in diversification rates during the Neogene, spurred by the Andean uplift, whereas spatial variation in Paleogene diversification rates may have left a weaker imprint on the current spatial patterns of palm and bird richness.

In addition to the time window considered, the contribution of spatial variation in net diversification rates to current patterns of species richness is likely to vary with spatial grain size and extent (Rahbek and Graves 2001). Spatial variation in species richness measured using small grain sizes across small spatial extents is likely determined mainly by the local influence of immigration and extinction (Rosenzweig 1995). The net sum of speciation and extinction events is thought to become increasingly important as spatial grain size (Rahbek and Graves 2001; Rosenzweig 1995) and extent (Evans et al. 2005) increase, assuming speciation occurs most often within relatively large areas dissected by dispersal barriers (Coyne and Orr 2004; Kisel and Barraclough 2010). Thus, the relative contribution of diversification rate to spatial patterns of species richness is expected to increase with spatial grain size and extent. Accordingly, evidence supporting the DRH often comes from studies focusing on richness patterns measured using relatively large spatial grain sizes across global or continental extents (e.g. Cardillo et al. 2005; Jablonski et al. 2006; Pyron 2014; Qian et al. 2007; Ricklefs 2006c; Ricklefs et al. 2006). The contribution of net diversification rates to richness patterns at finer spatial scales within continents remains largely unexamined (but see Jetz et al. 2012; Svenning et al. 2008).

Here, we first describe a novel approach to assess the explanatory power of the DRH by addressing the two quantitative questions posed above. Accordingly, we start by formally defining for any given time (window for diversification) and spatial scale scale: (1) the contribution of net diversification to current species richness in an area, and (2) the amount of spatial variance in current species richness that can be statistically explained by the spatial pattern of net diversification rate. Then we describe estimators of these two quantities, which can be obtained for any taxon and region given data on the current spatial pattern of species richness and a dated phylogeny. We show that these estimators are biased: they represent the maximum possible contribution of net diversification to current species richness in an area and the maximum amount of spatial variance in current species richness that can be statistically explained by the spatial pattern of net diversification rate, respectively. Therefore, these estimators quantify the maximum possible explanatory power of the DRH, and of any related hypothesis proposing that current spatial patterns of species richness are explained by the effect of environmental factors (such as energy availability, topographic heterogeneity, biotic interactions or climatic oscillations) on diversification rates. It follows that low estimate values would reject the DRH as a major explanation of spatial patterns of species richness. Thus, despite their bias, the estimators we present can be used to empirically challenge the DRH.

After introducing the approach, we present an empirical application using data on the spatial pattern of species richness of the Furnariidae, a large radiation of Neotropical birds across South America in conjunction with a time-calibrated phylogeny. In particular, we use various spatial grain sizes and several time windows to measure the contribution of net diversification to spatial variation in species richness. We show that the DRH may potentially explain much of the spatial variance in species richness as measured over time windows extending ≥15 Ma into the past, but only minor portions of the spatial variance over the last 10 Ma and measured at grain sizes commonly used in biogeography and macroecology (1° × 1°). Thus, our results illustrate how our approach can expose severe limitations in the explanatory power of the DRH.

## METHODS

### Definitions

We focus on addressing the two quantitative questions about the DRH posed in the introduction: (1) what is the contribution of net diversification over a given time period to current species richness in an area?, and (2) how much of the spatial variance in current species richness can be explained by the spatial pattern of net diversification rate? In this section we formally define the quantities central to these questions.

#### Contribution of net diversification to current species richness in an area

We start by defining the contribution of net diversification from some past time (hereafter *T*) until the present to the species richness of a group of organisms in any given region. Current species richness in a spatial sampling unit *i* (*species_i_*) is determined by species richness existing in that spatial unit at time *T*, and by the sum of speciation, immigration, and extinction events since time *T* (Jablonski et al. 2006; Rosenzweig 1995): 

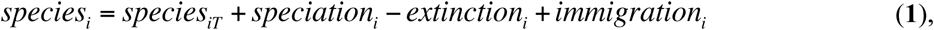
where *species_iT_* is the number of species that occurred in sampling unit *i* at time *T; speciation_i_* is the number of speciation events that took place since time *T* and were completely contained within the sampling unit *i* or partly overlapped the sampling unit *i; immigration_i_* is the number of immigration events into sampling unit *i* since time *T;* and *extinction_i_* is the number of extinction events within sampling unit *i* since time *T*. In the right-hand side of equation 1 species richness at time *T* is added to the three types of events that determine the gain and loss of species, thus describing the net accumulation (or loss) of species in spatial sampling unit *i* since time *T* until the present. Note that a single speciation event may overlap several sampling units and, therefore, may increase species richness in various sampling units (Fig. S1). Thus, our approach does not assume that speciation events can be referred uniquely to any particular sampling unit. We also stress that *extinction_i_* encompasses both global and local extinction. In the latter case, species disappear from sampling unit *i* but still occur outside it. Therefore, *extinction_i_* and *immigration_i_* account for local changes in species richness owing to shifts of species’ geographic distributions (Jablonski et al. 2006).

Based on equation 1, we defined the contribution of net diversification after time *T* to species richness in sampling unit *i* as:

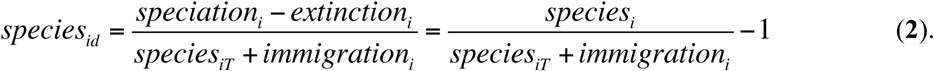

The quantity *species_id_* is the increase of *species_i_* due to net diversification since time *T*. As such, it is the net balance of extinction and speciation events occurring (fully or partially) within sampling unit *i*, divided by the initial number of species that had the potential to undergo speciation or extinction in sampling unit *i* after time *T*. This initial number of species includes not only all species present in sampling unit *i* at time *T* (i.e., *species_iT_*), but also all species that immigrated into sampling unit *i* after *T* (i.e., *immigration_i_*). Because *species_id_* is expressed as a proportion of the initial number of species, it can be thought of as a measure of the proportional increase of *species_i_* due to diversification, analogous to parameters measuring proportional increase in the number of individuals in population biology (Gotelli 2008). The proportional increase of *species_i_* due to diversification could, in principle, be defined over several segments of the period spanning from time *T* to the present. However, here we analyze only the net result from time *T* to the present, as expressed in equation 2. Note that *species_id_* can be negative, zero or positive.

#### Spatial variance in species richness explained by net diversification rate

Based on equations 1 and 2, species richness in a sampling unit *i* can now be expressed in terms of *species_id_*:

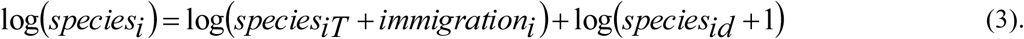

We used a logarithmic scale in equation 3 because diversification is a multiplicative process. The last term in equation 3 represents the contribution of *species_id_* to species richness in sampling unit *i*. Next we partition the spatial variance in species richness (i.e., the variance in log(*species_i_*) among sampling units) into three additive components according to equation 3 and properties of the variance of sums (Adler 2013):

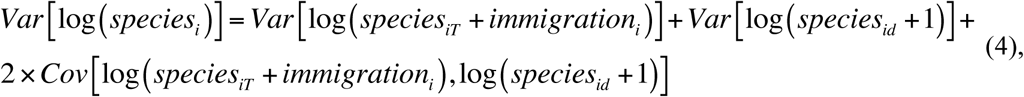
where 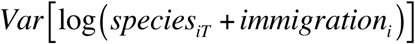 is the spatial variance across the study area of the initial number of species that had the potential to undergo speciation or extinction after time *T;* 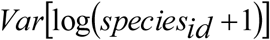 is the contribution of spatial variation in net diversification after time *T* (*species_id_*) to the current spatial pattern of species richness; and 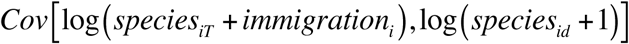 is the covariance of the previous two terms. The second term in the right-hand side of equation 4 is the amount of spatial variance in current species richness that is statistically explained by the spatial pattern of net diversification rate. This amount can be expressed as the ratio of spatial variance in species richness to spatial variance in *species_id_*: *Var[Log(species_id_* +*1)] / Var[Log(species_i_)]*. Note, however, that this ratio can be greater than 1 when the covariance term in equation 4 is negative.

#### Estimators

Unbiased estimation of the two quantities defined above *(species_id_* and *Var[Log (species_id_ +1)] / Var[Log(species_i_)])* is unlikely in most contexts because the values of *species_iT_* and *immigration_i_* in equation 2 are unknown; a possible exception are study systems with an outstanding fossil record (e.g., Jablonski et al. 2006). Next we describe estimators for both quantities that explicitly acknowledge this limitation and, yet, allow quantification of the maximum possible value of *species_id_* and *Var[Log (species_id_* +*1)] / Var[Log(species_i_)]*. We use carets to denote estimates of parameter values and thus distinguish them from the true parameter values. For instance, *speĉies_id_* denotes an estimate of *species_id_*.

#### Contribution of net diversification to current species richness in an area

We used the following approximation to estimate *species_id_*:

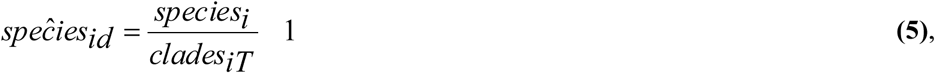
where *clades_iT_* is the number of clades that (a) originated from branches on a phylogenetic tree that intersect time *T* and (b) currently occur in sampling unit *i* (Fig. 1). Approximating the sum of *species_iT_* and *immigration_i_* (in equation 2) to *clades_iT_* (in equation 5) introduces bias in the estimate of *species_id_*. In particular, there are two (and only two) differences between *species_id_* and *speĉies_id_*: unlike *species_id_*, *speĉies_id_* assumes (1) that *immigration_i_* = 0 and (2) that none of the species that occurred in sampling unit *i* at time *T* went extinct leaving no descendants in sampling unit *i*, so that *species_iT_* = *clades_iT_*. If these two assumptions are met, then *species_i_* would always be greater than or equal to *clades_iT_*, and *clades_iT_* would equal *species_iT_* + *immigration_i_*. Therefore, in contrast to *species_id_*, *speĉies_id_* can only be zero (when *species_i_* = *clades_iT_*) or positive (when *species_i_* > *clades_iT_*), never negative. Moreover, the two assumptions leading to differences between *species_id_* and *speĉies_id_* correspond to two scenarios under which *speĉiesid* overestimates *speĉies_id_*. First, if *immigration_i_* > 0, then *species_iT_* + *immigration_i_* ≥ *clades_iT_* and thus *speĉies_id_* could overestimate *species_id_*. Second, if any of the species that occurred in sampling unit *i* at time *T* went extinct leaving no descendants in sampling unit *i*, then *species_iT_* + *immigration_i_* > *clades_iT_* and thus *speĉies_id_* would overestimate *species_id_*. It thus follows that *speĉies_id_* ≥ *species_id_*. In other words, the direction of the bias involved in our approach is known: when *speĉes_id_* is not equal to the contribution of net diversification after time *T* to species richness in sampling unit *i*, it is always an overestimate of this quantity. Note also that because fewer diversification events are included in the calculation of *speĉies_id_* as *T* is closer to the present, *speĉies_id_* cannot increase with decreasing *T* and it must decline or remain constant as *T* decreases.

**Figure 1.**
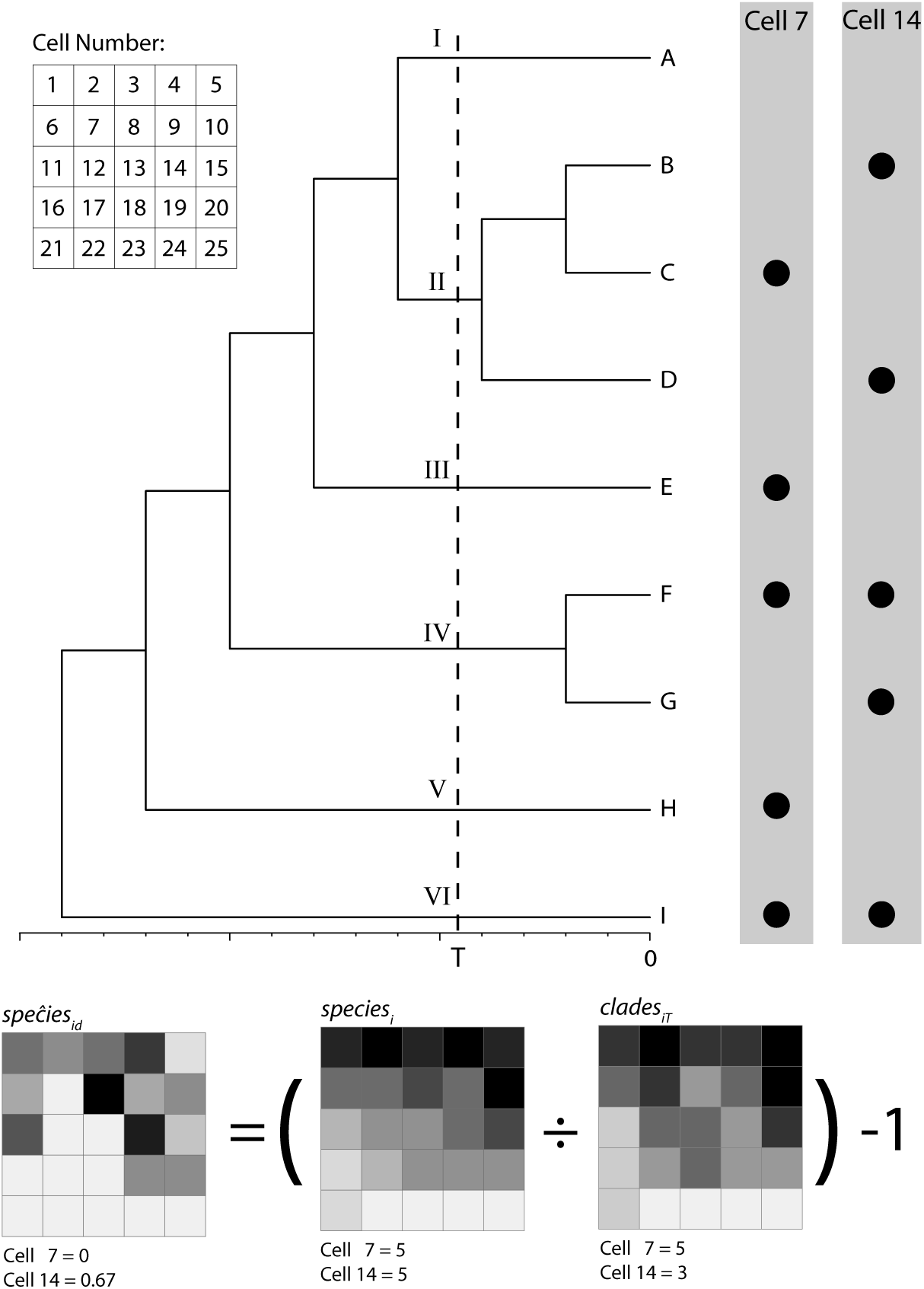
Illustration of spatial sampling units with equal species richness and, yet, different contributions of diversification after time *T* to local species richness. Extant species occurring in sampling units 7 and 14 are shown as dots to the right of the phylogenetic tree, and maps below show *speĉies_id_*, i.e., the contribution of net diversification to current species richness in each unit (see equation 5 in the text). All lineages leading to the species occurring in sampling unit 7 diverged from one another before time *T;* thus, diversification occurring after time *T* did not contribute to current species richness in this sampling unit and *speĉies_id_* equals zero whereas *clades_iT_* equals five. In contrast, two pairs of species currently occurring in sampling unit 14 diverged after time *T* (species B and D, and species F and G); here, diversification after time *T* did contribute to current species richness. Because *speĉies_id_* equals 0.667 in this case, one can conclude that diversification after time *T* increased diversity in this sampling unit by at most 66.7%. The relatively high value of *speĉies_id_* in sampling unit 14 could also reflect immigration events resulting in co-occurrence of species that diverged after time *T*, and not only the direct effect of net diversification on the species richness of sampling unit 14. Note that *speĉies_id_* does not equal the average net diversification rate (NDR) of the clades present in a sampling unit: because species C does not occur in sampling unit 14, *speĉies_id_* is not equal to the average NDR of clades II, IV, and VI.

#### Spatial variance in species richness explained by net diversification rate

One can not directly estimate 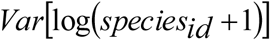 because unbiased estimates of *species_id_* for each sampling unit are lacking. Because *speĉes_id_* is likely an overestimate of *species_id_* (see previous paragraph), for sampling unit *i* one can be certain that *species_id_* is somewhere in the interval between zero and *speĉies_id_*. This interval information does not allow one to calculate the variance, but it is sufficient to calculate the maximum possible value of the variance (Dharmadhikari and Joagdev 1989; Ferson et al. 2007). Accordingly, we calculated the maximum possible value of 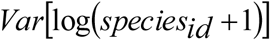, hereafter 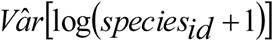, using two approaches. First, following Dharmadhikari and Joagdev (1989) we calculated it as:

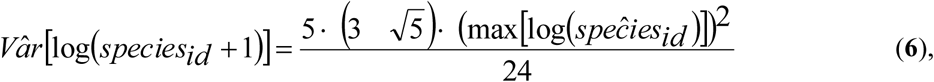
where 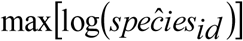 is the maximum value of 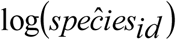 across the study region. This first approach assumes that the cumulative distribution function of *species_id_* is *S*-shaped, i.e., that it crosses the uniform cumulative distribution function once from below. This assumption is less stringent than the assumption of unimodality: it is met by all unimodal distributions and by some multimodal distributions (Dharmadhikari and Joagdev 1989). Second, we calculated 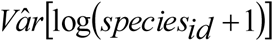 using the computational approach described in section 4.5.1.4 in Ferson et al. (2007). This latter approach has no distributional assumptions; it produces estimates of the maximum possible value of the variance that are certain to enclose the true variance, but they are conservative in that they may be higher than the exact maximum value (Ferson et al. 2007).

Having calculated the maximum possible value of the contribution of spatial variation in net diversification after time *T* to the current spatial pattern of species richness, one can now estimate the maximum possible value of the ratio of spatial variance in *species_id_* to the overall spatial variance in species richness: 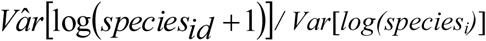. We now turn to an empirical application of our approach.

### Study system and analysis

We focus on South America as a domain for our analyses because the continent contains regions with unusually high species richness (Myers et al. 2000; Orme et al. 2005), mainly caused by rapidly diversifying groups that evolved in isolation within the boundaries of this biogeographic domain (Fjeldså 1994; Fjeldså and Irestedt 2009; Fjeldså and Rahbek 2006; Mittelbach et al. 2007; Rahbek et al. 2007; Rahbek and Graves 2001; Schemske 2009). Several clades show marked spatial variation in species richness within South America. One such clade comprises the ovenbirds and woodcreepers (Passeriformes, familly Furnariidae), an endemic radiation of passerine birds with over 250 species occurring in all terrestrial environments in the continent. Spatial patterns of diversity of furnariids resemble those of the overall avifauna, with diversity peaking in topographically complex areas in the tropical Andes and in warm and humid zones in the Amazon region (see below). These characteristics make furnariids an appropriate clade to examine the contribution of diversification rates to spatial patterns of species richness.

Our analyses are based on the maximum clade credibility (MCC) tree of the Furnariidae (*sensu* Remsen et al. 2015) reported by Derryberry et al. (2011), which corresponds to Scleruridae, Dendrocolaptidae, Xenopsidae and Furnariidae in the classification of Ohlson et al. (2013). This chronogram was constructed using a number of geological calibration points (see Derryberry et al. 2011). We defined a total of 170 values of *T*, from 34 Ma to the present with intervals of 0.2 Ma, to examine the extent to which diversification after various points in time along the evolutionary history of furnariids could explain current spatial patterns of species richness. Although some of our main conclusions rely on relative, not absolute times, we address possible biases resulting from uncertainty in the temporal calibration of the phylogenetic tree in the Discussion.

We based our geographical analyses on distribution maps of the 269 South American furnariid species sampled in Derryberry et al. (2011). These maps are the product of extensive review of museum collections and literature, comprehensive fieldwork, and interpolation based on consideration of variation in habitat and species-habitat associations (Fjeldså and Irestedt 2009; Rahbek et al. 2007; Rahbek and Graves 2001). These maps consist of a grid with a total of 1489 1°×1° sampling units with > 90 % land coverage where each species is coded as present or absent.

To examine the influence of spatial grain size in our results, we quantified furnariid species richness across South America using ten grain sizes from 1°×1° to 10°×10°, based on the geographical distribution of species mapped at a resolution of 1°×1°. For each grain size we subsequently estimated, for the 165 time intervals defined above, (1) the maximum contribution of net diversification to current species richness in an area (i.e., *speĉies_id_*), and (2) the maximum spatial variance in species richness attributable to net diversification rate (i.e., 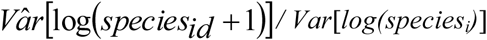).

Both approaches to estimating the maximum proportion of the variance in species richness attributable to net diversification rate (see section on estimators above) produced similar trends relative to the time intervals over which net diversification rates were measured. However, for small grain sizes the approach based on Ferson et al. (2007) yielded lower values than that based on Dharmadhikari and Joagdev (1989), despite the fact that the former approach may yield overestimates. This difference suggests that the assumptions required by the approach based on Dharmadhikari and Joagdev (1989) were not met. Thus, for small grain size, we focus on results produced by following Ferson et al. (2007) because they do not rely on assumptions about the distribution of *speĉies_id_*.

## RESULTS

### Contribution of net diversification to current species richness in an area

We found that for all grain sizes the maximum contribution of net diversification to species richness on each sampling unit (*speĉies_id_*) decreased with the value *T* defining the time window considered (Figs. 2 and 3). Although a smaller value of *speĉies_id_* is expected when *T* approaches the present (see section on estimators above), this decrease was not constant and was particularly steep in between time intervals defined by the deeeper nodes in the Furnariidae phylogeny. Net diversification during the last 33 Ma may have increased the number of species in an average 1°×1° area by a factor of >13, and may have done so by a factor > 37 in an average 10°×10° area. In contrast, net diversification over the last 10 Ma did not even double the number of species in most of the areas considered here (Figs. 2 and 3).

**Figure 2.**
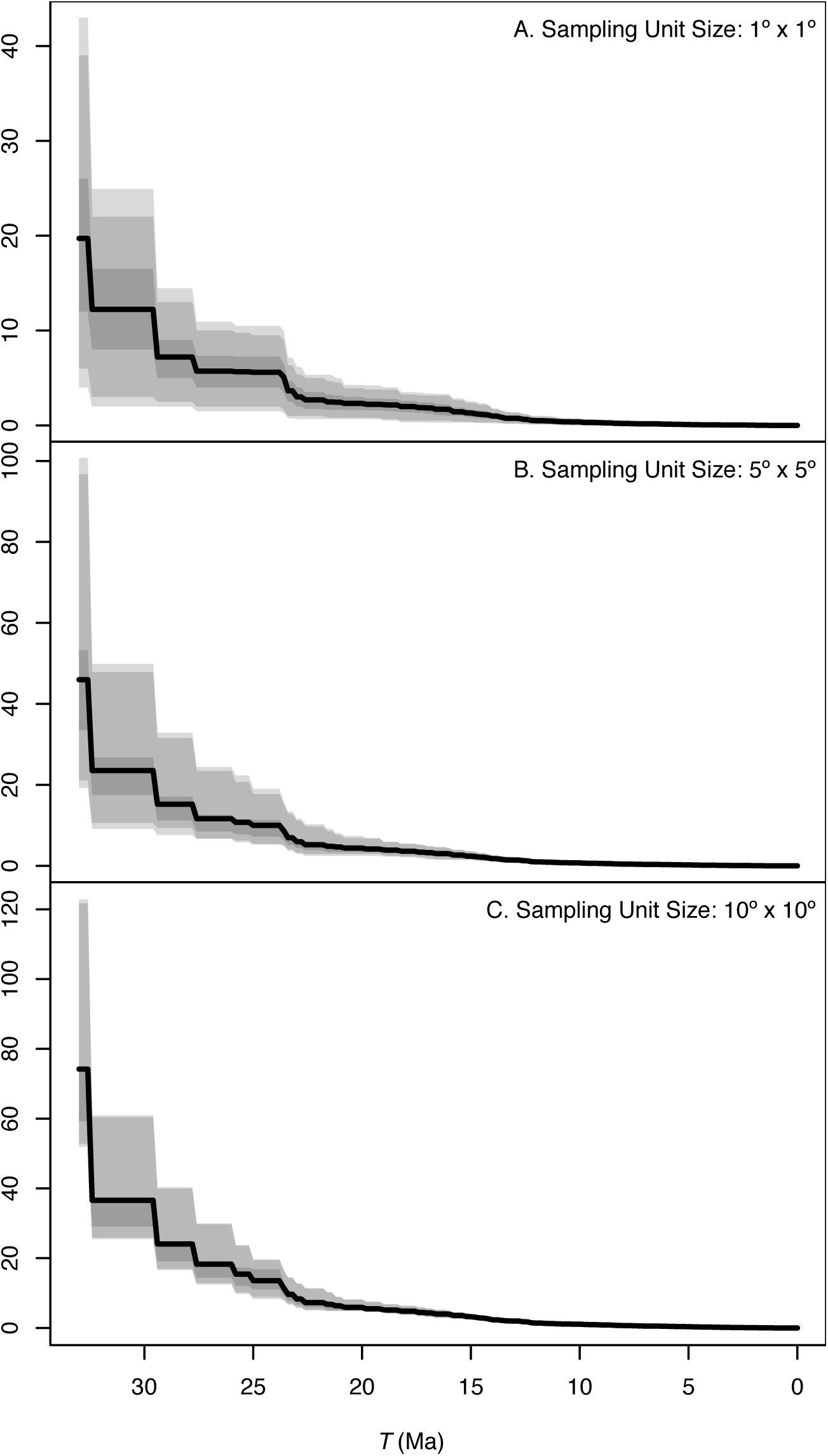
Relationship between the value of *T* and the maximum contribution of net diversification to furnarid species richness in a sampling unit (*speĉies_id_*) for three grain sizes across South America: (A) 1° × 1°, (B) 5° × 5° and (C) 10° × 10°. The solid lines represent the mean value of *speĉies_id_* across the study area and shades indicate the 75, 90 and 95 percentiles for any given value of *T* in the abscissa. Note that the scale of the ordinate differs among panels; the magnitude of and variation in *speĉies_id_* is lower for smaller grain sizes.

**Figure 3.**
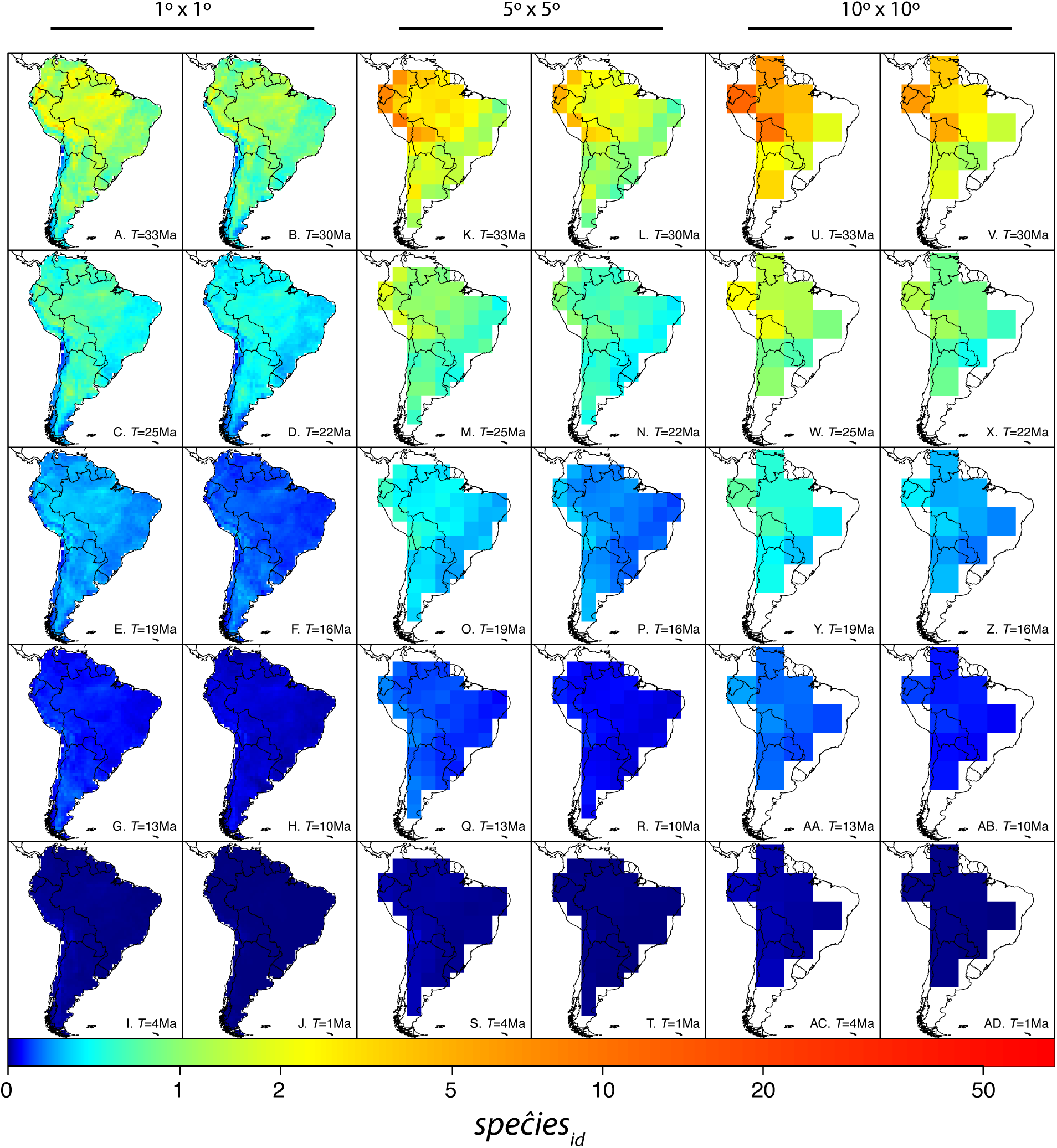
Spatial variation of the maximum contribution of net diversification to fumariid species richness in an area (*speĉies_id_*) for different values of *T* (from 33Ma to 1Ma) and grain sizes: (A-J) 1° × 1°, (K-T) 5° × 5° and (U-AD) 10° × 10°. The color scale at the bottom applies to all maps.

The maximum contribution of net diversification to species richness was not uniform across the continent. For example, while net diversification during the past 33 Ma multiplied the number of species in sampling units in the tropical Andes by a maximum factor of 12 to 20 (depending of grain size), it only multiplied species richness in sampling units in Eastern Brazil or the Southern Andes by a maximum factor of 2 to 9 in the same time period (Fig. 3A, 3K and 3U). Noticeable spatial variation in *species_id_*, at least for *T ≥* 25 Ma (Fig. 3), suggests that spatially variable net diversification rates may explain spatial variation in current species richness. We examine this possibility next.

### Spatial variance in species richness explained by net diversification rate

The maximum fraction of the spatial variance in species richness accounted by spatial variation in net diversification rates also declined for all grain sizes as *T* was closer to the present (Fig. 4). However, for the period corresponding to the first half of the furnariid radiation (i.e., *T ≥* 15.2 Ma), all spatial variance in species richness could be accounted for by variation in diversification rate (Fig 4). The time interval (from *T* to the present) in which not all the spatial variance in current species richness could be accounted for by the variance in net diversification rate was shorter when larger grain sizes were considered (15 Ma in 1° × 1°, 7.5 Ma in 10° × 10°; Fig 4). For values of *T* when only a fraction of the spatial variance in current species richness could be accounted for by variance in *species_id_* (i.e., *T*< 15.2 Ma), diversification tended to account for a lower proportion of variance in species richness at smaller grain sizes (Fig. 4). Indeed, the more reliable approach to estimating the spatial variance in species richness explained by net diversification rate (i.e., the computational approach of Ferson et al. (2007); see methods) indicated that the DRH can explain, at most, < 16% of this variance for *T ≤* 10 Ma and < 2.5% for *T ≤* 5 Ma (Fig. 4B). Thus, these results uncover stern upper limits to the explanatory power of the DRH, as applied to diversification events taking place since the Late Miocene and to one of the most commonly used spatial scales to characterize species richness patterns in biogeography and macroecology (1° × 1°).

**Figure 4.**
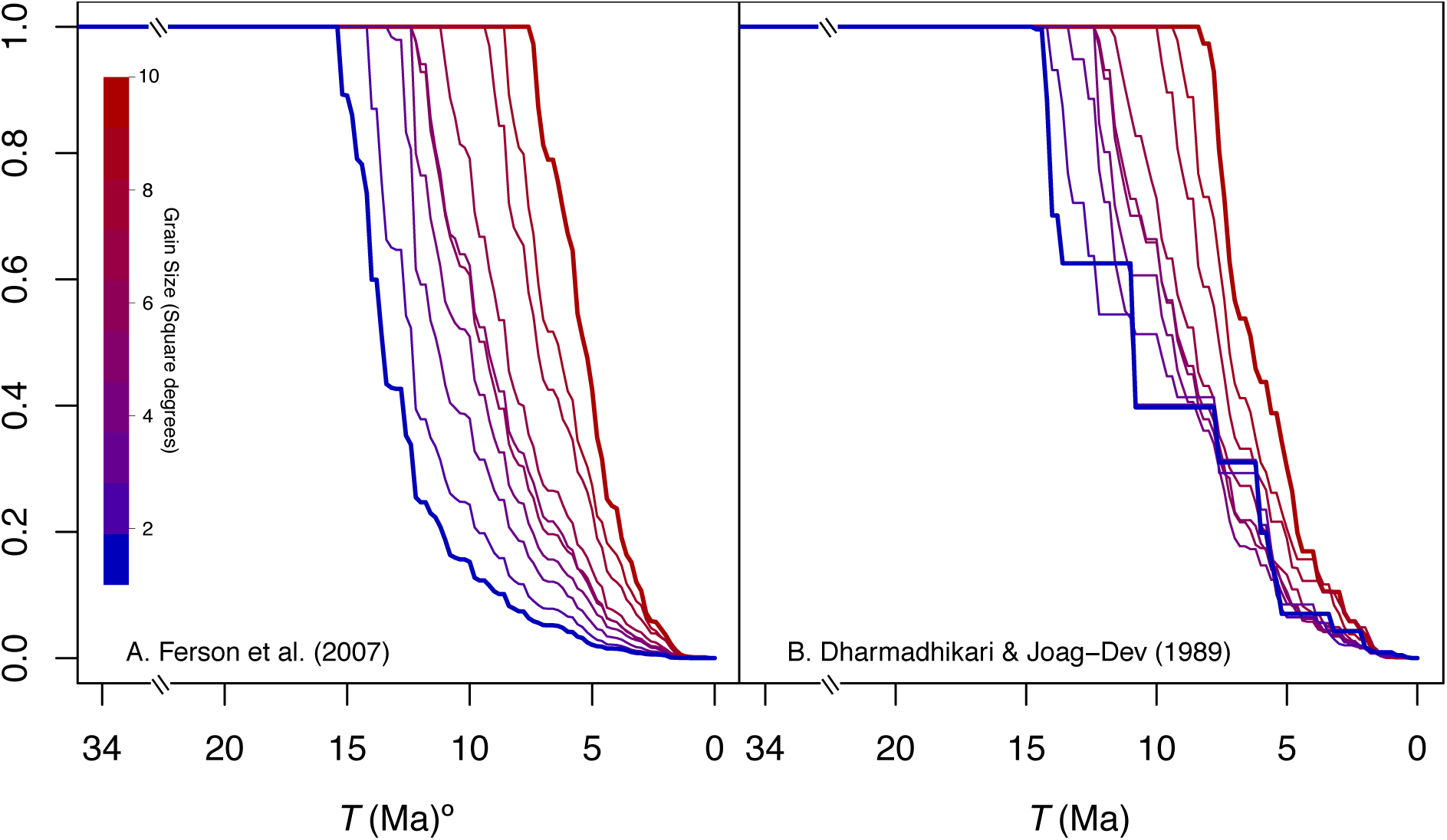
Maximum proportion of the variance in species richness across spatial sampling units that can be explained by diversification, measured for various time windows extending from time *T* to the present. Lines present values for different spatial grain sizes colored from blue to red (1° × 1° to 10° × 10°, respectively). The upper bound of the variance in *species_id_* across spatial sampling units is estimated using (A) the approach described in Ferson et al. (2007) which has no distributional assumptions and (B) following Dharmadhikari and Joagdev (1989), assuming that the cumulative distribution function of *species_id_* is *S*-shaped. For values of *T* where only a fraction of the spatial variance in current species richness could be accounted for by variance in *species_id_*, the maximum possible explanatory power of the DRH increases with spatial grain size.

## DISCUSSION

The diversification rate hypothesis (DRH) proposes that current spatial patterns of species richness result from variation in net diversification rates (Fischer 1960; Mittelbach et al. 2007; Ricklefs 2004). Here we developed an approach to address two questions central to understanding the explanatory power of the DRH: (1) what is the contribution of net diversification over a given time period to current species richness in an area?, and (2) how much of the spatial variance in current species richness can be explained by the spatial pattern of net diversification rate? We applied the approach to study geographical variation in species richness of funariid birds across South America, and found the answers to the two questions above to be contingent on the value of *T* defining the time window over which net diversification rate was measured and on the spatial grain size used to measure species richness. Regarding the first question, for all grain sizes, the maximum contribution of net diversification rate to species richness in an area declined with *T*, and ranged from 13 – 37 fold local increases in species richness for *T* = 33 Ma to less than 2-fold increases for *T ≤* 10 Ma. Regarding the second question, the maximum proportion of spatial variance in current species richness explained by spatial variation in net diversification rate also declined with *T* for all grain sizes. Spatial variation in net diversification rate could potentially account for all spatial variance in species richness when *T ≥* 15.2 Ma regardless of grain size. For smaller values of *T*, spatial variation in net diversification rate could explain less variance in species richness over small than over large grain sizes. At the smallest grain size we considered (1° × 1°), diversification rates could only explain < 16% of the variance for *T ≤* 10 Ma and < 2.5% for *T* ≤ 5 Ma. Taken together, these findings allow only for a very limited explanatory power of the DRH as applied to diversification events occurring since the Late Miocene and at 1° × 1° grain size, a very common spatial scale in studies of species richness in biogeography and macroecology.

It is difficult to overemphasize the role that the definitions we used (see Definitions in Methods) play in addressing the two quantitative questions on which our study focuses. In particular, equation 1 describes how the species richness of any given area arises from three fundamental processes: speciation, immigration, and extinction. We stress that this equation does not imply that speciation always occurs within any single area, such as a 1° × 1° or 10° × 10° sampling unit. In other words, equation 1 does not assume that speciation necessarily occurs at the spatial scale at which the richness pattern is measured. Equation 1 simply formalizes the uncontroversial claim that a speciation event can potentially increase species richness in any single sampling unit. This indeed happens when a speciation event is completely contained within a sampling unit, but also (and arguably more often, especially so at the smaller spatial grain we examined) when a speciation event partly overlaps a sampling unit (Fig. S1). Similarly, equation 1 does not assume that extinction necessarily occurs at the spatial scale at which the richness pattern is measured. The extinction term in equation 1 includes cases in which a species disappears from a sampling unit but still occurs outside that sampling unit (i.e., local extirpation, in some cases caused by range shifts), as well as cases of global extinction.

The approach we presented here is based on the assumption that it may often be virtually impossible to answer accurately our two focal questions. This is because describing geographic details of speciation, immigration, and extinction events that took place in the distant past is not feasible given that the geographic distributions of species and clades may shift substantially over time. In terms of our definitions, this means that the values of *species_iT_* and *immigration_i_* in equation 2 are unknown, unless an exceptional fossil record is available (e.g. Jablonski et al. 2006). Accordingly, we focused on developing estimators that provide biased answers to our two focal questions. Crucially, however, we have demonstrated the direction of the bias is known, and therefore the estimators yield meaningful answers to our study questions. In particular, these estimators yield maximum possible values for (1) the contribution of net diversification over a given time period to current species richness in an area and (2) the spatial variance in current species richness that can be explained by the spatial pattern of net diversification rate. The bias in estimators needs to be considered by any empirical study adopting our approach, as we illustrate in the remainder of the discussion.

The interpretation of our results must be appraised by considering the potential influence of immigration on patterns of species richness. Range dynamics leading to immigration into sampling units, as well as differential persistence of lineages outside and inside such units, may affect our estimate of the contribution of spatial variation of diversification rates to the spatial variance in species richness. As previously noted, *speĉies_id_* likely overestimates the contribution of diversification to current species richness in sampling unit *i* because it is also affected by immigration events resulting in co-occurrence within sampling unit *i* of species that diverged after time *T*. The likelihood of this happening increases with *T* because the probability of change in geographic distributions leading to range overlap increases with time (Losos and Glor 2003). Thus, time windows defined by larger values of *T* would suffer from a more severe upward bias in *speĉies_id_* as an estimate of *species_id_*. Therefore, the observed decline with *T* in *speĉies_id_* and in the estimated proportion of spatial variance in current species richness explained by spatial variation in net diversification rate 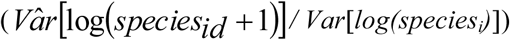 may partly reflect differences in the magnitude of bias introduced by immigration. Nevertheless, we emphasize that while we cannot empirically assess the influence of immigration, the estimates we report here remain meaningful as maximum possible values of *species_id_* and *Var[Log (species_id_ +1)] / Var[Log(species_i_)]*(see methods). Therefore, our approach places upper limits on the explanatory power of the DRH even if one cannot accurately estimate such power. Because our results reveal that the maximum explanatory power of the DRH is reduced over recent times and small spatial grains, regardless of the unknown effects of immigration we can safely conclude that spatial variation in net diversification rates over recent times is not a sufficient explanation of spatial patterns of species richness, which are often the focus of biogeographic and macroecological analyses.

The increased impact of diversification rate on the spatial richness pattern with increasing grain size is consistent with the idea that speciation likely occurs most often within relatively large areas that allow for geographic separation of populations (Coyne and Orr 2004; Kisel and Barraclough 2010; Losos and Schluter 2000). Therefore, one would expect that as the size of spatial sampling units (i.e., grain size) increases, sampling units slide along a continuum from “island-ness” to “province-ness” (sensu Rosenzweig 1995), such that species richness is increasingly dominated by the balance between speciation and extinction events (i.e., *species_id_)* at increasingly larger scales (Fig 2). Alternatively, because immigration rates may be higher in larger than smaller spatial units (Rosenzweig 1995), it is possible that our estimates of the contribution of diversification to spatial variance in species richness are more strongly biased for larger than for smaller grain sizes owing to the unmeasured effect of immigration on *speĉies_id_*. Again, these estimates are nonetheless valid as maximum possible values, and the following discussion emphasizes their implications as such.

The link between diversification and spatial patterns of species richness can arise when localized phenomena influence net diversification rate only within particular regions, including the emergence of barriers promoting allopatric speciation, the occurrence of catastrophic events promoting extinction, or the colonization of new areas or environments where lineages may diversify (Goldberg et al. 2011; Moore and Donoghue 2007). In these cases diversification would be structured spatially and would contribute to spatial variation in species richness. For example, our analyses suggest that net diversification prior to 15 Ma could contribute substantially to variance in current species richness as measured with a grain size of 1°×1° latitude-longitude sampling units (Fig. 4). This potential effect of net diversification on species richness would have been largely restricted to the northern Andes, the Venezuelan Guyana, and regions within Amazonia depending on grain size (Figs. 3A–F, 3K–P, 3U–Z). The potential impact of net diversification on species richness in the Andes and the Venezuelan Guyana may have been particularly strong because topographic heterogeneity may foster allopatric speciation as populations persist and adapt to stable local environments (Fjeldså 1994; Fjeldsâ et al. 2012; Fjeldså and Rahbek 2006; Rahbek et al. 2007; Rahbek and Graves 2001). In contrast, a particularly high contribution of net diversification to species richness in some regions of Amazonia may reflect the persistence of small relictual clades of mid-Tertiary age within warm-humid tropical biomes, likely remnants of the formerly more extensive tropics (Fjeldså and Irestedt 2009).

The most significant implication of the empirical findings in this study is the limited ability of the DRH to explain richness patterns of furnariids, measured at small grain sizes, by resorting to diversification events that occurred since the Late Miocene. By example, whereas diversification over the past 5 Ma contributed importantly to the overall number of species in furnariids (Brumfield 2012), spatial variation in diversification rate over this time period can account for at most < 2.5% of the spatial variance in current species richness among 1°×1° areas. This implies that at a small-grain scale, recent diversification in furnariids had a rather geographically uniform (i.e., not spatially structured) effect on species richness patterns. This apparent mismatch between the time when the clade diversified extensively and the time when diversification rate contributed most to fine-scale spatial variance in current patterns of richness might reflect fairly homogeneous birth-death process of clade growth (Brumfield 2012; Derryberry et al. 2011) or pulses in diversification driven by the evolution of key traits or by reasons such as global climate change affecting diversification regardless of the geographic context (Moore and Donoghue 2007). In these scenarios, overall richness may have increased with no important effect on spatial variance in current species richness. In any case, furnariid diversification during the Plio-Pleistocene (last 5 Ma) has contributed notably to the overall extant diversity of the clade (Brumfield 2012), but cannot explain much of the spatial variance in current species richness at the smaller grain sizes commonly used in continental-wide studies of species richness. This may appear surprising in the light of studies like that of Weir (2006), which indicated that during times as recent as the Pleistocene, birds in the Neotropical highlands experienced high diversification rates, whereas those in the lowlands did not. If such spatial variation in diversification rates were indeed a general phenomenon, then our analyses suggest that it contributed significantly only to the spatial patterns of species richness when measured at large spatial scales rarely considered in macroecological studies (but see Rahbek and Graves 2001).

Likewise, although the uplift of the Andes undoubtedly had an impact on diversification by creating opportunities for geographic isolation and for adaptation to new environments since the Late Miocene (Gregory-Wodzicki 2000), such processes appear not to have substantially influenced spatial patterns of richness, as measured using small grain sizes, through their effects on net diversification of furnariids. Furnariid net diversification during the last 10 Ma could explain at most <16% of the spatial variance in species richness. Therefore, hypotheses alternative to the DRH may best explain how Andean uplift during relatively recent times might have affected spatial richness patterns. Some of these hypotheses emphasize the role of immigration as a determinant of richness in any given area (see equation 1). By example, new environments created by the uplift of the Andes may have increased richness mainly by fostering immigration of species, largely from lineages originating in the southern cone of the continent that were pre-adapted to cool or open environments (see Chapman 1917; Fjeldså and Irestedt 2009; Kennedy et al. 2014). Such an ecological “fitting” or “sorting” process (Agosta and Klemens 2008) may have been particularly important in determining the diversity of various Andean groups including birds (Chapman 1917) and plants (Donoghue 2008).

Our finding that the DRH is unable to explain spatial patterns of diversity at small grain sizes by invoking diversification events since the Late Miocene relies heavily on the time frame of furnariid diversification implied by molecular phylogenetic analyses being accurate (Derryberry et al. 2011). If the dates in the phylogeny were positively biased (i.e., node age-estimates are older than they actually are), then the explanatory power of the DRH would be even more limited than we calculated. Alternatively, if estimates of dates in the phylogeny were negatively biased (node-age estimates are younger than they actually are), then we would be underestimating the explanatory power of the DRH. More generally, even if the empirical results in our particular study system are compromised by uncertainty in the temporal calibration of the phylogeny, the approach we present to estimate the explanatory power of the DRH would still be valid, and the empirical study would at the very least show how the DRH could potentially be challenged based on a hypothetical example.

In conclusion, the approach we developed here allows evaluating the maximum explanatory power of the DRH at different spatial scales of analysis and considering different time intervals. Our results indicate that the DRH may play a major role in explaining current species richness patterns over large scales because spatial and temporal variation in diversification rates may have left a long-term signal in contemporary spatial patterns of furnariid species richness beyond playing a major role in establishing the overall source pool of species. Indeed, to the extent that contemporary environmental factors vary among sampling units with varying species richness, and considering that our analyses revealed that diversification may account for considerable fractions of spatial variance in richness over large scales, our work echoes suggestions that diversity-environment relationships may arise from large-scale evolutionary processes (Kozak and Wiens 2012; Pyron and Wiens 2013; Ricklefs 2006a). However, our empirical findings for furnariids also indicate that the role of the DRH as a major explanation is, at the very least, restricted to richness patterns measured at large grain sizes, or to diversification rates measured before the Late Miocene. Thus, our study uncovered important upper limits to the explanatory power of the DRH and, in that way, contributes insight into the relative contribution of various factors to determining geographical patterns of species richness.

The generality of the results reported here needs to be assessed by analyses of organisms with different histories, such as groups in which recent colonization of South America was followed by rapid diversification including not only birds (Barker et al. 2015) but also other organisms like plants (Hughes and Eastwood 2006) and mammals (Webb 2006). More generally, we expect that additional studies estimating the contribution of diversification to current spatial patterns of diversity will improve our understanding of how speciation, extinction and immigration act in concert to shape the distribution of biological diversity.

## Supporting Information

**Figure S1.**
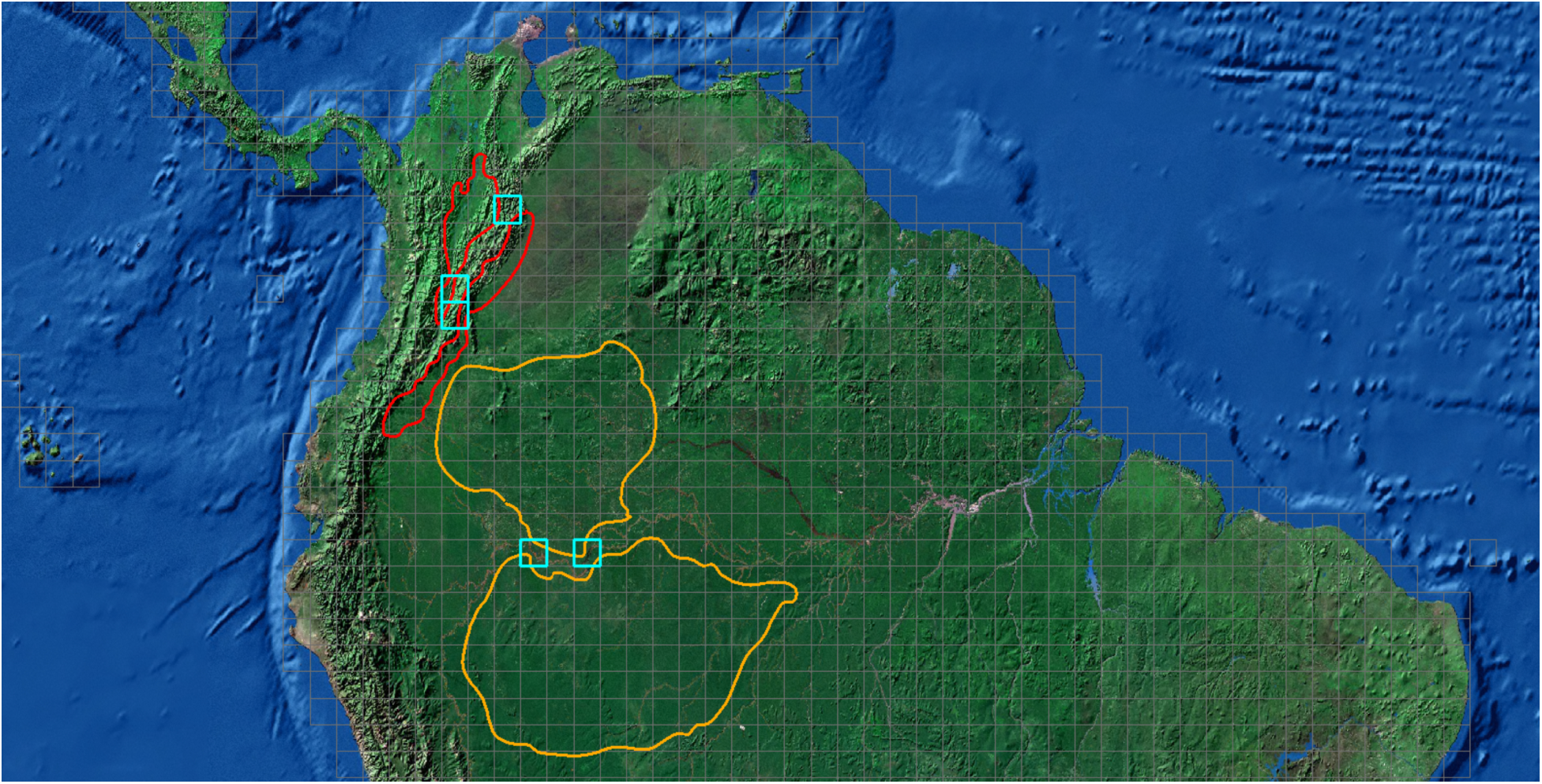
Hypothetical example showing two speciation events that increase species richness in more than one sampling unit. Sampling units are 100 km × 100 km squares defined by the gray grid. The polygons show the distribution of two pairs of sister species (orange and red) at the time of the speciation events. This speciation events increase richness in two and three sampling units respectively, outlined in cyan.

